# Adaptation of Iterative Ultrastructure Expansion Microscopy (iU-ExM) enables high-resolution imaging of the cytoskeleton and subcellular architecture of *Trypanosoma cruzi*

**DOI:** 10.64898/2026.10.28.762540

**Authors:** Normanda Souza-Melo, Giovanna Henriques de Souza, Sharmila Ortiz, Wanderley de Souza

## Abstract

Kinetoplastid parasites, including *Trypanosoma cruzi*, possess a highly organized subpellicular microtubule network essential for maintaining cell shape, organelle positioning, and dividing. Here we describe results obtained using immunofluorescence microscopy of cells processed using conventional ultrastructure expansion microscopy (U-ExM), that improved visualization of the cytoskeletal structures, with an ∼4.5-fold expansion, as well as the recently described and iterative ultrastructure expansion microscopy (iU-ExM) with about 6.2-fold and 18-fold expansion of epimastigotes of *T. cruzi*. The optimized protocol involves the formation of an initial hydrogel crosslinked with N,N′-(1,2-dihydroxyethylene) bisacrylamide (DHEBA), a temperature-and alkaline pH-sensitive cleavable crosslinker, followed by thermal denaturation at 85 °C, intermediate immunostaining, neutral gel re-embedding, and the formation of a second hydrogel crosslinked with N,N′-methylenebisacrylamide (BIS).

Subsequent dissolution of the first hydrogel under alkaline conditions enables a second round of expansion. This approach achieved approximately 6.2-fold expansion during the first round and a final linear expansion of approximately 18-fold. Using conventional confocal microscopy, iU-ExM enabled high-resolution visualization of individual subpellicular microtubules and specialized internal microtubule arrangements, including those associated with the basal body, flagellar pocket, Golgi complex, and cytopharynx. Furthermore, the enhanced spatial resolution facilitated the examination of cytoskeletal reorganization at different stages of the cell cycle, revealing changes in microtubule organization associated with mitotic spindle formation and cytokinesis. Together, these findings establish iU-ExM as an accessible high-resolution imaging approach for investigating the complex subcellular architecture of *T. cruzi*, providing new opportunities to explore cytoskeletal organization and remodeling during parasite cell division using conventional confocal microscopy.

## Research Highlights

We adapted iU-ExM to *Trypanosoma cruzi*, achieving approximately 18x expansion. Confocal microscopy resolved specialized microtubule arrays and revealed cytoskeletal organization during mitosis and cytokinesis.

## Graphical Abstract

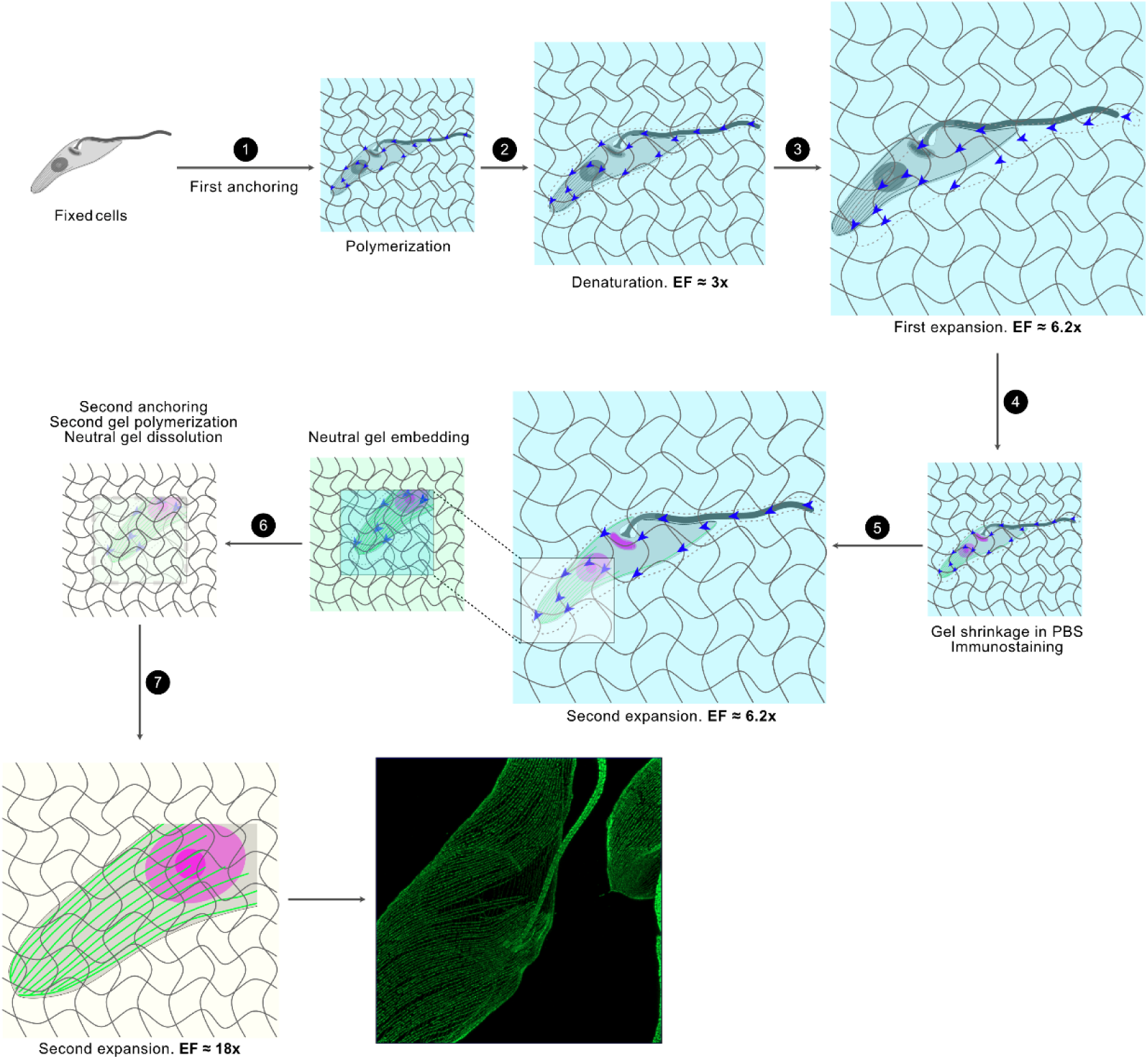

## 1. Introduction

Protozoan parasites belonging to the family Trypanosomatidae have been extensively investigated using a wide range of microscopic approaches aimed at understanding their structural and functional organization. These organisms are of considerable biological and medical interest due to their complex life cycles, which involve distinct morphological forms and, in many species, transitions between vertebrate and invertebrate hosts. Their cellular architecture is characterized by several distinctive features, including the kinetoplast, a specialized region of the mitochondrion containing a highly organized network of mitochondrial DNA composed of minicircles and maxicircles, typically located near the flagellar basal body. Another remarkable feature is the presence of a highly organized network of subpellicular microtubules positioned immediately beneath the plasma membrane, forming a corset-like structure that contributes to cell shape, polarity, and the spatial organization of intracellular components (Alcantara et al., 2025; de Souza, 1984, 2009; Gull, 1999; Vidal & Souza, 2017; Zuma et al., 2021). Beyond their unique cellular organization, several members of this family, including *T. cruzi*, *Trypanosoma brucei*, and *Leishmania* spp., are responsible for parasitic diseases of major human and veterinary importance, such as Chagas disease, human African trypanosomiasis, and leishmaniasis (de Souza, 2009; de Souza et al., 2026; Gull, 1999).

The cytoskeleton of trypanosomatids exhibits a remarkable degree of structural specialization, comprising not only the subpellicular microtubule network but also distinct microtubule assemblies associated with the flagellum, basal bodies, flagellar pocket, and other intracellular compartments (Alcantara et al., 2025; de Souza, 2009; de Souza et al., 2026; Lacomble et al., 2009; Sant’Anna et al., 2005; Vidal & Souza, 2017). In *T. cruzi*, the cytostome–cytopharynx complex represents a particularly specialized endocytic structure supported by two distinct sets of microtubules: a quartet originating near the flagellar pocket and a triplet associated with the cytostome. These microtubules accompany the cytopharynx and contribute to its characteristic three-dimensional organization (Alcantara et al., 2014; Vidal et al., 2016). The spatial relationships among the flagellar apparatus, specialized microtubule arrays, and intracellular organelles highlight the complexity of kinetoplastid cellular architecture (de Souza, 2009; Lacomble et al., 2009, 2012; Rotureau et al., 2013). Importantly, these cytoskeletal structures undergo coordinated remodeling during the cell cycle. In *T. cruzi*, the cytostome–cytopharynx complex disassembles during cell division, whereas its associated microtubule quartet persists in a shortened form and duplicates during the G2/M transition, contributing to the subsequent reassembly of the complex in daughter cells (Alcantara et al., 2017). Understanding the three-dimensional organization and remodeling of these specialized microtubule arrangements is therefore essential for elucidating the mechanisms underlying parasite morphogenesis, organelle organization, cell division, and differentiation (Alcantara et al., 2017; Vidal & De Souza, 2017, 2022).

Over the past decades, electron microscopy has played a fundamental role in revealing the ultrastructural organization of trypanosomatids. A broad range of approaches, from conventional transmission electron microscopy (TEM) of ultrathin sections to electron microscopy tomography and cryo-electron tomography, has provided detailed information about the spatial relationships among cytoskeletal elements and intracellular organelles (Alcantara et al., 2014; Gluenz et al., 2015; Lacomble et al., 2009; Sant’Anna et al., 2005). However, these techniques generally require sophisticated instrumentation, specialized sample preparation, and extensive image acquisition and processing, particularly for three-dimensional reconstructions. Moreover, the analysis of large cell populations and the identification of specific molecular components can be challenging. In contrast, fluorescence microscopy enables the examination of numerous cells and the localization of selected proteins using specific antibodies or fluorescent markers. Advances in confocal and light super-resolution microscopy have substantially improved the investigation of cellular architecture. Nevertheless, conventional fluorescence microscopy remains constrained by the diffraction limit, with a lateral resolution of approximately 200–250 nm, preventing the discrimination of individual subpellicular microtubules and other closely associated subcellular structures (Chen et al., 2015; Gorilak et al., 2021).

Expansion microscopy (ExM) has emerged as an alternative approach to overcome these limitations by physically enlarging biological specimens rather than relying exclusively on improvements in optical resolution (Chen et al., 2015). This technique involves anchoring cellular components to a swellable polymer network that expands upon hydration, thereby increasing the physical distances between structures that would otherwise remain unresolved. Subsequent methodological advances led to ultrastructure expansion microscopy (U-ExM), which combines optimized sample anchoring, hydrogel embedding, and thermal denaturation to improve the preservation and visualization of cellular ultrastructure (Chen et al., 2015; Gambarotto et al., 2019). Conventional U-ExM typically achieves approximately 4–4.5-fold linear expansion and has been successfully applied to kinetoplastid parasites. (Gorilak et al., 2021) demonstrated its applicability to *T. brucei* and *Leishmania major*, enabling quantitative analyses of cytoskeletal organization. Subsequently, Alonso et al., 2022 applied U-ExM to different developmental stages of *T. cruzi*, allowing the visualization of microtubule organization and the localization of tubulin isoforms and isotypes. Beyond kinetoplastids, expansion microscopy has also improved the visualization of cytoskeletal and subcellular structures in other protozoan parasites, including *Plasmodium* spp. and *Giardia lamblia* (Bertiaux et al., 2021; Jetishi et al., 2024). In trichomonads, conventional ExM enabled the visualization of individual axostyle microtubules by fluorescence microscopy (Bandeira et al., 2023). Despite these advances, the expansion factors achieved with conventional U-ExM remain limited for resolving densely packed microtubule arrays and specialized internal cytoskeletal structures.

More recently, iterative ultrastructure expansion microscopy (iU-ExM) was developed to achieve substantially greater physical expansion through successive rounds of hydrogel embedding and physical expansion (Alonso, 2022; Louvel et al., 2023). This approach combines the ultrastructural preservation provided by U-ExM with an iterative expansion strategy involving an initial hydrogel crosslinked with N,N′-(1,2-dihydroxyethylene)bisacrylamide (DHEBA), a cleavable crosslinker sensitive to temperature and alkaline pH. Following thermal homogenization and intermediate immunolabeling, the specimen is re-embedded in a neutral hydrogel and subsequently incorporated into a second expansion hydrogel crosslinked with N,N′-methylenebisacrylamide (BIS). The initial DHEBA-based network is then dissolved under alkaline conditions, enabling a second round of physical expansion. This strategy can achieve approximately 16–20-fold linear expansion, substantially improving the effective spatial resolution attainable with conventional fluorescence microscopes. Given the highly organized and complex cytoskeletal architecture of *T. cruzi*, iU-ExM represents a promising approach for investigating the three-dimensional organization of its subcellular structures, particularly the densely packed subpellicular microtubule network and specialized internal microtubule assemblies.

In the present study, we describe the adaptation and optimization of iU-ExM for high-resolution imaging of the cytoskeleton and subcellular architecture of *T. cruzi*. By implementing a sequential hydrogel expansion strategy, we achieved a final linear expansion factor of approximately 18-fold, enabling the visualization of individual subpellicular microtubules and specialized internal microtubule arrangements using conventional confocal fluorescence microscopy. This approach also allowed the three-dimensional examination of cytoskeletal organization and its structural remodeling at different stages of the cell cycle, particularly during mitosis and cytokinesis. Our findings demonstrate the potential of iU-ExM as a valuable methodological tool for investigating the complex cellular architecture of *T. cruzi* and provide a foundation for future studies of cytoskeletal organization and parasite cell division.

## 2. Materials and Methods

### 2.1. Parasite culture

Epimastigote forms of *Trypanosoma cruzi* (Dm28c wild-type strain) were cultivated in liver infusion tryptose (LIT) medium supplemented with 10% fetal bovine serum (FBS), 10 U/mL penicillin, and 10 μg/mL streptomycin. Cultures were maintained at 28 °C in the exponential growth phase through regular subculturing, with parasites seeded at a density of 5 x 10⁶ cells/mL.

### 2.2. Iterative ultrastructure expansion microscopy (iU-ExM)

#### 2.2.1. Parasite preparation and fixation

Approximately 1×10^7^ epimastigotes were washed with phosphate-buffered saline (PBS) and resuspended in 300 µL of PBS. The parasite suspension was transferred onto 13-mm round glass coverslips previously coated with poly-L-lysine and placed in 24-well plates. Parasites were allowed to adhere to the coverslips for 10 min at room temperature. Subsequently, the supernatant was removed, and the adhered parasites were washed with PBS and fixed with 500 µL of a solution containing 0.1% glutaraldehyde (GA) and 3% paraformaldehyde (PFA) in PBS for 15 min at room temperature. Following fixation, the fixative solution was removed, and the samples were washed three times with PBS. The coverslips were then immediately processed for expansion microscopy.

#### 2.2.2. Gelation chamber assembly

iU-ExM was performed based on the protocol described by Louvel et al. 2023, with modifications for *Trypanosoma cruzi*. Briefly, a humidified chamber was prepared using a Petri dish containing filter paper moistened with water (Supplementary Figure S1). For each gelation chamber, a piece of Parafilm measuring approximately 60 x 30 mm was placed over the moistened filter paper. The gelation chamber was assembled directly on the Parafilm using two stacks of glass coverslips, each consisting of two 18 x 18 mm coverslips with an individual thickness of 0.17 mm. The coverslips within each stack were held together using approximately 5 µL of water, and excess water was carefully removed with filter paper. The two stacks were positioned parallel to each other, approximately 9 mm apart, to accommodate a 13-mm round coverslip containing the fixed parasites. The parasite-containing coverslip was then positioned between the two stacks, with the adhered parasites facing the Parafilm, creating a confined space for hydrogel polymerization with an approximate thickness of 340 µm.

#### 2.2.3. First anchoring and hydrogel polymerization

The fixed parasites on glass coverslips were incubated with 500 µL of an anchoring solution containing 2% acrylamide (AA – Bio-Rad) and 1.4% PFA in PBS for 3 h at 37 °C. Following anchoring, excess solution was carefully removed using absorbent tissue, and the coverslips were positioned in the previously assembled gelation chambers (Section 2.2.2). The monomer solution, containing 10% AA, 19% sodium acrylate (SA – AK Scientific), and 0.1% N,N′-(1,2-dihydroxyethylene) bisacrylamide (DHEBA - Sigma), was prepared one day before use, without the polymerization initiators, and stored at −20 °C. The solution was divided into 95 µL aliquots to allow individual thawing immediately before use. To prevent premature polymerization, ammonium persulfate (APS – Bio-Rad) and tetramethylethylenediamine (TEMED – Bio-Rad) were added only immediately before gelation, at final concentrations of 0.25% each. The gelation chambers were maintained in a humidified environment, and the activated monomer solution was added to completely fill the space surrounding the parasite-containing coverslips. The samples were incubated on ice for 15 min, followed by incubation at 37 °C for 45 min to complete hydrogel polymerization.

#### 2.2.4. Denaturation and first expansion

Following hydrogel polymerization, the gelation chamber was inverted onto a flat surface, and the coverslip containing the polymerized gel was carefully removed from the chamber. The coverslip was then transferred, with the gel facing upward, to a 6-well plate containing 2 mL of denaturation buffer (200 mM SDS, 200 mM NaCl, and 50 mM Tris base, pH 6.8). The samples were maintained under gentle agitation until the gel detached from the coverslip. The detached gel was subsequently transferred to a 1.5 mL microcentrifuge tube containing 1 mL of fresh denaturation buffer and incubated at 85 °C for 90 min. The temperature was carefully monitored throughout the incubation. Following denaturation, the gel was transferred to a Petri dish (90 x 15 mm) containing ultrapure water, ensuring that the gel was fully submerged. The water was replaced at least three times to promote hydrogel swelling until a stable linear expansion factor of approximately 6.2x was achieved. The gel could also remain submerged in ultrapure water overnight to ensure that swelling reached equilibrium.

#### 2.2.5. Intermediate immunolabeling

Following the first expansion, ultrapure water was gradually replaced with PBS until the gel contracted to approximately 3 cm in size. The gel was then transferred onto a clean surface covered with Parafilm and carefully cut into approximately 1 x 1 cm fragments. The gel fragments were either stored in 24-well plates containing 500 µL of PBS supplemented with 0.01% sodium azide at 4 °C or immediately processed for immunolabeling. For immunolabeling, gel fragments were transferred to a 24-well plate containing 500 µL of blocking buffer (3% bovine serum albumin [BSA] in PBS) and incubated for 3 h at room temperature. The gels were subsequently washed 3 times for 10 min each with washing buffer (PBS containing 0.1% Tween-20). The samples were then incubated overnight at room temperature with 300 µL of mouse monoclonal anti-TAT-1 antibody (Woods et al., 2013; kindly provided by Prof. Keith Gull), diluted 1:400 in blocking buffer supplemented with 0.01% sodium azide. Following primary antibody incubation, the gels were washed 3 times for 10 min each with washing buffer and subsequently incubated with 300 µL of Alexa Fluor 488-conjugated anti-mouse IgG secondary antibody (Invitrogen), diluted 1:400, for 3 h at room temperature. The gels were then washed 3 times for 10 min each with washing buffer.

For DNA staining, the gels were incubated with 300 µL of Hoechst 33342 (Sigma) solution (40 µg/mL in PBS) for 1 h, followed by three washes of 20 min each with washing buffer. Following immunolabeling, the gels were immersed in ultrapure water and allowed to expand until their dimensions reached a stable plateau. Prior to imaging, 35-mm glass-bottom dishes with a 14-mm microwell and a No. 1.5 coverglass (0.16–0.19 mm thickness) were treated with 0.1% poly-L-lysine for 10 min at 37 °C and subsequently rinsed with water. The expanded gels were then transferred to the treated dishes and imaged using a ZEISS LSM 900 confocal laser scanning microscope equipped with 20x, 40x, and 63x objectives.

#### 2.2.6. Second expansion - Neutral gel embedding and second anchoring

Following intermediate immunolabeling, the gel was expanded and cut into a 1 x 1 cm fragment. The expanded gel fragment was transferred to a 6-well plate and incubated 3 times for 10 min each on ice, under gentle agitation, in 2 mL of a freshly prepared neutral gel solution containing 10% AA, 0.05% DHEBA, 0.1% APS, and 0.1% TEMED in ultrapure water. Following incubation, the gel was placed on a glass microscope slide inside a humidified chamber consisting of a Petri dish containing filter paper moistened with water. Two folded aluminum foil spacers, adjusted to match the thickness of the gel, were positioned on either side of the gel fragment. The gel was placed between the spacers and covered with a 22 x 22 mm glass coverslip. The assembly was then incubated at 37 °C for 1 h to allow polymerization of the neutral gel.

Following polymerization, the gel was transferred to a 6-well plate and subjected to a second anchoring step by incubation in 2 mL of a solution containing 2% AA and 1.4% PFA in ultrapure water for 3 h at 37 °C under gentle agitation. Subsequently, the gel was washed with PBS for 30 min.

#### 2.2.7. Second expansion - Second gel polymerization and final expansion

For the second expansion, the gel was incubated three times for 10 min each on ice, under gentle agitation, in 2 mL of a previously prepared second expansion monomer solution containing 10% AA, 19% SA, 0.1% N,N′-methylenebisacrylamide (BIS), 0.1% APS, and 0.1% TEMED. Excess monomer solution was carefully removed using filter paper. The gel was then transferred onto a glass microscope slide inside a humidified chamber. Two folded aluminum foil spacers were positioned on either side of the gel to support a 22 x 22 mm glass coverslip. The assembly was incubated at 37 °C for 1 h to allow polymerization of the second expansion gel.

Following polymerization, the gel was transferred to a 6-well plate and incubated in 3 mL of 200 mM NaOH for 1 h at room temperature under gentle agitation to dissolve the first expansion gel and the neutral gel. The gel was subsequently washed three times for 15 min each with PBS, until the pH of the washing solution reached approximately 7. Finally, the gel was transferred to distilled water, which was replaced several times until the gel dimensions reached a stable plateau, corresponding to a final linear expansion factor of approximately 18x.

For confocal imaging, the fully expanded gel was initially examined using a 20x objective to identify the side containing the parasite samples. The gel was then carefully cut horizontally into two thinner sections using a scalpel blade to reduce its thickness and facilitate imaging with higher-magnification objectives. The section containing the samples was transferred to a poly-L-lysine-coated glass-bottom dish, prepared as previously described, and imaged using a ZEISS LSM 900 confocal laser scanning microscope equipped with 40x and 63x objectives. The samples were subsequently imaged using a ZEISS LSM 900 confocal laser scanning microscope equipped with 40x and 63x objectives.

#### 2.2.8. Confocal imaging, image processing, and expansion factor determination

Following expansion, the hydrogels were mounted in 35-mm glass-bottom dishes with a 14-mm microwell and a No. 1.5 coverglass (0.16–0.19 mm thickness), previously coated with 0.1% poly-L-lysine, as described above. For imaging after the second expansion, the hydrogels were initially examined using a 20x objective to identify the side containing the parasite samples. The gels were then carefully sectioned horizontally using a scalpel blade to reduce their thickness and facilitate high-magnification imaging.

Confocal imaging was performed using a ZEISS LSM 900 confocal laser scanning microscope equipped with 20x, 40x, and 63x objectives. Image processing and visualization were performed using ZEN software (ZEISS) and Fiji (ImageJ). Final figures were assembled using Affinity Designer.

The linear expansion factor was determined by comparing the dimensions of the hydrogels before and after expansion. The final expansion factor (EF) achieved after the second expansion was approximately 18x. EF = Expanded gel diameter (mm)/13 (mm).

## 3. Results

### 3.1. Adaptation of the iU-ExM workflow for *Trypanosoma cruzi*

To overcome the resolution limitations of conventional fluorescence microscopy and standard ultrastructure expansion microscopy (U-ExM), we adapted iterative ultrastructure expansion microscopy (iU-ExM) for the analysis of the cytoskeletal architecture of *T. cruzi* epimastigotes (Figure 1). This adaptation incorporated a sequential hydrogel expansion strategy based on two polymer networks with distinct crosslinking properties. The first network, containing the cleavable crosslinker N,N′-(1,2-dihydroxyethylene) bisacrylamide (DHEBA), provided the basis for the initial expansion, whereas the second, crosslinked with N,N′-methylenebisacrylamide (BIS), supported the subsequent expansion cycle. Thermal denaturation, intermediate immunolabeling, neutral gel re-embedding, and alkaline cleavage of the DHEBA-containing network were integrated into this strategy, enabling successive physical enlargement of the parasite specimens while retaining fluorescently labeled cytoskeletal structures.

**Figure 1:**
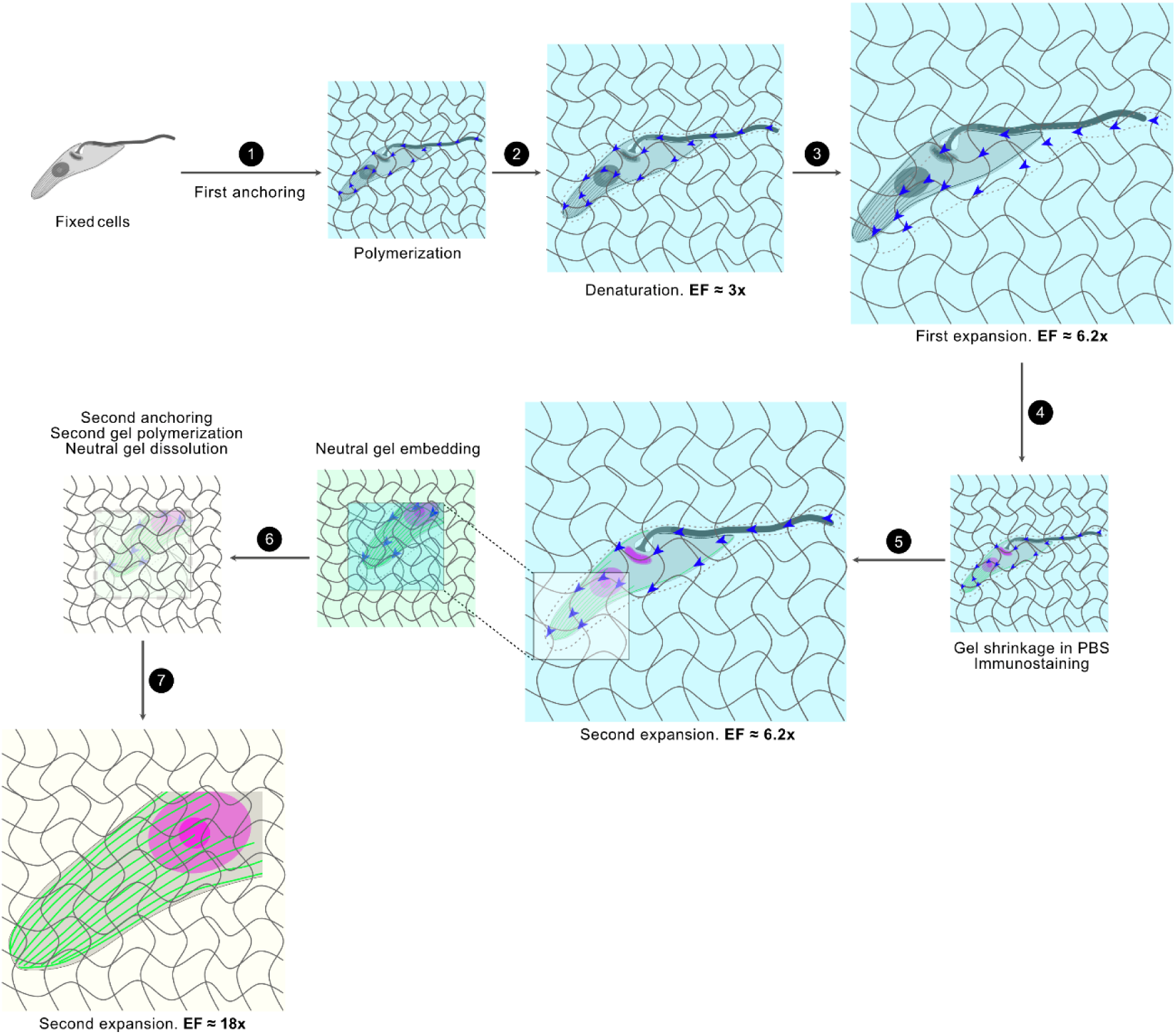
Schematic representation of the iterative ultrastructure expansion microscopy (iU-ExM) workflow adapted for *Trypanosoma cruzi*. The protocol consists of two successive rounds of hydrogel embedding and expansion, with intermediate immunolabeling to preserve fluorescence labeling throughout the iterative process. (1) Fixed parasites are anchored to a first expandable hydrogel containing acrylamide, sodium acrylate, and the cleavable crosslinker N,N′-(1,2-dihydroxyethylene)bisacrylamide (DHEBA). (2) The embedded specimens undergo thermal denaturation at 85 °C for 1h 30 min, followed by initial hydrogel expansion in water to promote physical separation of cellular structures. (3) The first expansion is completed through successive water exchanges, reaching a linear expansion factor of approximately 6.2x. (4) The expanded hydrogel is contracted in PBS to facilitate sample handling and immunolabeling with anti-TAT-1 antibody, followed by fluorescent secondary antibody and Hoechst 33342 staining. (5) The immunolabeled specimen is re-expanded in water and embedded in a neutral hydrogel to preserve the spatial organization of the labeled structures during subsequent processing. (6) Following a second anchoring step, the specimen is embedded in a second expandable hydrogel containing N,N′-methylenebisacrylamide (BIS) as a non-cleavable crosslinker. The intermediate hydrogel is subsequently dissolved by incubation in 200 mM NaOH. (7) The specimen undergoes final expansion in water, reaching an overall linear expansion factor of approximately 18x and enabling high-resolution visualization of the parasite’s cytoskeletal architecture by confocal fluorescence microscopy.

The adapted iU-ExM approach achieved an initial linear expansion factor of approximately 6.2x, which increased to approximately 18x following the second expansion cycle. This substantial enlargement increased the physical separation between neighboring cytoskeletal elements, facilitating their discrimination by conventional confocal microscopy. Importantly, the expanded epimastigotes retained identifiable morphological features and recognizable microtubule arrangements, enabling the examination of cytoskeletal structures within the context of the overall parasite architecture. The combination of successive expansion and intermediate immunolabeling thus provided a means of investigating the spatial organization of closely associated microtubule populations that are difficult to distinguish using conventional fluorescence microscopy. These results establish the feasibility of applying iU-ExM to *T. cruzi* and provide the methodological basis for the detailed characterization of its cytoskeletal architecture presented in the following sections.

### 3.2. Sequential hydrogel expansion during iU-ExM

To evaluate the physical expansion achieved during the adaptation of iU-ExM to *T. cruzi*, we monitored changes in hydrogel dimensions throughout the successive expansion steps (Figure 2). Following the first gel polymerization and thermal denaturation, the hydrogel exhibited an initial linear expansion of approximately 3x, increasing from an initial diameter of 13 mm to approximately 40 mm. Subsequent immersion in water promoted further swelling, resulting in a linear expansion factor of approximately 6.2x, corresponding to a gel diameter of approximately 80 mm.

**Figure 2:**
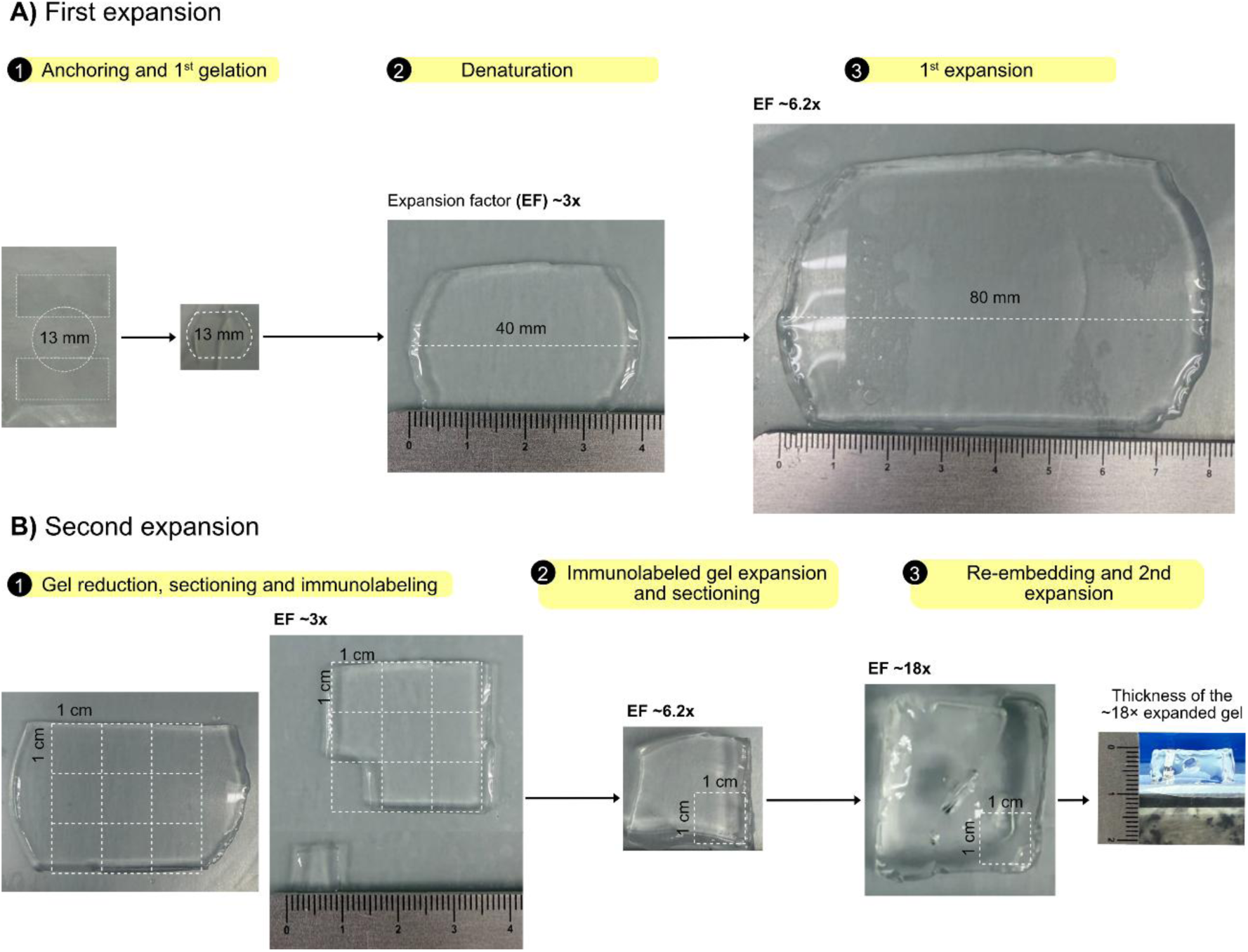
Sequential hydrogel expansion and sample preparation during iterative ultrastructure expansion microscopy (iU-ExM). Representative photographs illustrating the successive stages of hydrogel expansion and sample processing throughout the iU-ExM workflow adapted for *Trypanosoma cruzi*. A) Following the first anchoring and hydrogel polymerization (1), thermal denaturation at 85 °C for 90 min resulted in an initial linear expansion of approximately 3x (40 mm) (2), relative to the original 13-mm coverslip diameter. Subsequent immersion in water promoted further hydrogel swelling, reaching a linear expansion factor of approximately 6.2x (80 mm) (3). B) The expanded hydrogel was then contracted in PBS, sectioned into smaller fragments (1), and subjected to intermediate immunolabeling. Following immunolabeling, the hydrogel fragments were re-expanded in water and sectioned into approximately 1 x 1 cm pieces to facilitate subsequent processing. The fragments were then embedded in a neutral hydrogel, subjected to a second anchoring step, and incorporated into a second expandable hydrogel (2). After dissolution of the intermediate hydrogel and final expansion in water, an overall linear expansion factor of approximately 18x was achieved (3). Dashed lines indicate the measured dimensions of the hydrogels or representative regions selected for sectioning.

Following the first expansion, the hydrogel was reduced in size by replacing water with PBS, allowing the gel to be sectioned into smaller fragments for intermediate immunolabeling. After immunolabeling, the gel fragments were re-expanded in water and sectioned into approximately 1 x 1 cm fragments for subsequent re-embedding in a neutral hydrogel and incorporation into the second expandable matrix.

Following alkaline treatment to dissolve the DHEBA-containing gel networks, the second hydrogel underwent further swelling in water, reaching a final linear expansion factor of approximately 18x relative to the original sample dimensions. These observations demonstrate the progressive physical expansion achieved through successive hydrogel expansion cycles and establish the feasibility of implementing iU-ExM in *T. cruzi*.

### 3.3. First expansion reveals the cytoskeletal and subcellular architecture of ***Trypanosoma cruzi***

Following the first expansion, *T. cruzi* epimastigotes exhibited a linear expansion, enabling improved visualization of their cytoskeletal and subcellular organization by confocal microscopy (Figure 3). Immunolabeling with the anti-TAT-1 antibody, combined with Hoechst 33342 staining, allowed the identification of the subpellicular microtubule network, nucleus, and kinetoplast, while preserving the characteristic elongated morphology of the parasites (Figure 3A). Analysis of confocal z-stacks revealed distinct populations of internal microtubules. While projections encompassing a larger portion of the z-stack highlighted the overall parasite morphology and the subpellicular microtubule network (Figure 3B-C), projections generated from selected optical sections reduced signal overlap and improved the visualization of internal cytoskeletal structures.

**Figure 3:**
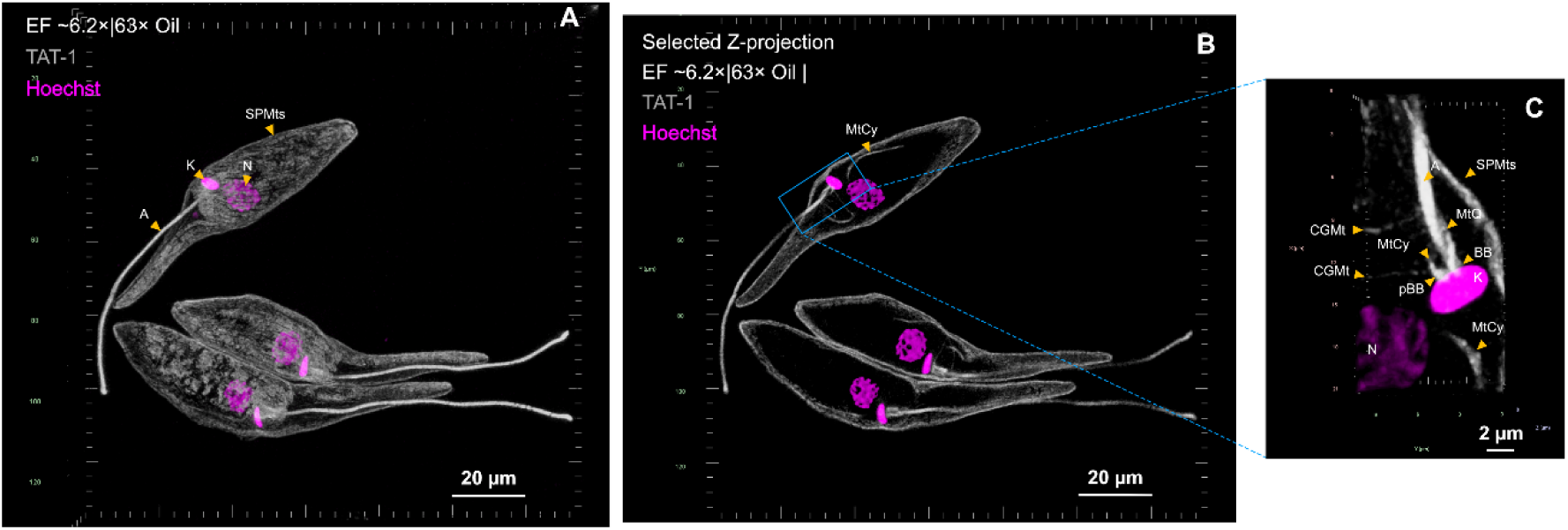
Visualization of the cytoskeletal and subcellular architecture of *Trypanosoma cruzi* epimastigotes following the first expansion. Representative confocal images of epimastigotes subjected to the first expansion cycle of iU-ExM, reaching a linear expansion factor of approximately 6.2x. Microtubules were labeled with the anti-TAT-1 antibody (gray), and DNA was stained with Hoechst 33342 (magenta). (A) Confocal z-stack projection showing the overall parasite morphology, including the subpellicular microtubule network (SPMTs), axoneme (A), nucleus (N), and kinetoplast (K). (B) Projection generated from selected optical sections of the z-stack, reducing the overlap of fluorescence signals from the subpellicular microtubule network and facilitating the visualization of internal cytoskeletal structures. The blue rectangle indicates the region enlarged in panel C. (C) Higher-magnification view highlighting specialized microtubule arrangements, including the microtubule quartet (MtQ), microtubules associated with the cytostome–cytopharynx complex (MtCy), and Golgi complex-associated microtubules (CGMt), as well as the basal body (BB), probasal body (pBB), axoneme (A), nucleus (N), and kinetoplast (K). Scale bars: 20 µm (A, B) and 2 µm (C), corresponding to expanded sample dimensions. All images were acquired using a 63x oil-immersion objective.

Detailed examination of the selected region revealed specialized microtubule arrangements associated with the flagellar pocket region, including the basal and probasal bodies, the microtubule quartet, and microtubules associated with the cytostome–cytopharynx complex and the Golgi complex (Figure 3C). These observations demonstrate that the first expansion provides sufficient spatial separation to distinguish specialized microtubule structures that are difficult to resolve by conventional fluorescence microscopy and whose ultrastructural organization in *T. cruzi* has historically been characterized primarily by transmission electron microscopy.

### 3.4. Iterative expansion enhances the visualization of subpellicular and internal microtubule architecture

Following the second expansion, *T. cruzi* epimastigotes exhibited a final linear expansion factor of approximately 18x, enabling a more detailed visualization of their cytoskeletal architecture by confocal microscopy (Figure 4). The increased physical separation between microtubules improved the visualization of the subpellicular microtubule network, revealing its highly ordered arrangement along the parasite cell body (Figure 4A).

**Figure 4:**
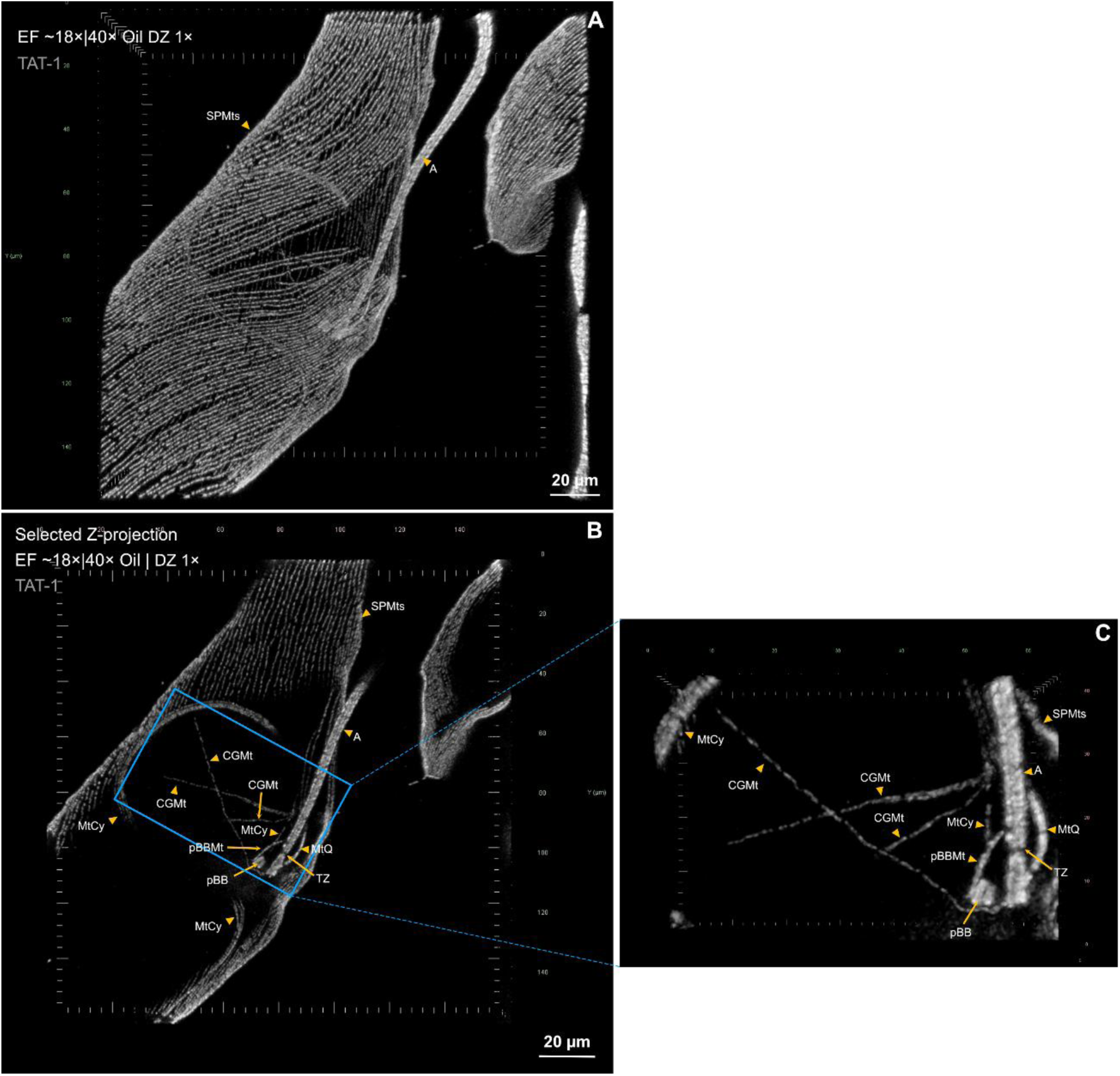
Iterative ultrastructure expansion microscopy reveals the subpellicular microtubule network and specialized internal microtubule arrays in *Trypanosoma cruzi*. Representative confocal images of epimastigotes subjected to iU-ExM, achieving a final linear expansion factor of approximately 18x. Microtubules were labeled with the anti-TAT-1 antibody and a fluorescent secondary antibody (gray). (A) Projection of the complete confocal z-stack showing the highly organized subpellicular microtubule network (SPMTs) and the axoneme (A). (B) Projection generated from selected optical sections of the z-stack, reducing the overlap of fluorescence signals from the subpellicular microtubules and allowing visualization of specialized internal microtubule arrays. The blue rectangle indicates the region enlarged in panel C. (C) Higher-magnification view highlighting the microtubule quartet (MtQ), cytostome–cytopharynx-associated microtubules (MtCy), Golgi complex-associated microtubules (CGMt), basal body (BB), probasal body (pBB), probasal body-associated microtubules (pBBMt) and transition zone (TZ), together with the axoneme (A) and subpellicular microtubules (SPMTs). Scale bars: 20 µm (A, B), corresponding to expanded sample dimensions. All images were acquired using a 40x oil-immersion objective with 1x digital zoom (DZ).

Analysis of the complete confocal z-stack highlighted the extensive distribution and organization of subpellicular microtubules throughout the parasite cell body, as well as a distinct region lacking subpellicular microtubules corresponding to the cytostome (Figure 4A). Three-dimensional visualization of the confocal image stack further illustrates the spatial arrangement and overall organization of this microtubule network (Supplementary Movie 1). In contrast, projections generated from selected optical sections reduced the overlap of fluorescence signals from the subpellicular microtubule network, allowing internal microtubule structures to be distinguished more clearly (Figure 4B).

Higher-magnification examination of the selected region revealed several specialized microtubule arrangements, including the microtubule quartet, microtubules associated with the cytostome–cytopharynx complex, and microtubules associated with the Golgi complex. The axoneme and additional structures associated with the flagellar apparatus, including the basal and probasal body regions and the transition zone, could also be distinguished (Figure 4C). Three-dimensional visualization of these structures provided additional spatial information, allowing the apparent points of origin and trajectories of individual microtubule arrays to be followed within the parasite (Supplementary Movie 2).

Together, these observations demonstrate that iterative expansion substantially improves the spatial separation and visualization of specialized microtubule populations in *T. cruzi*, enabling detailed examination of cytoskeletal structures that are difficult to distinguish after the first expansion.

### 3.5. iU-ExM reveals cytoskeletal remodeling during mitosis and cytokinesis in *Trypanosoma cruzi*

To further investigate cytoskeletal remodeling during the cell cycle, we examined *T. cruzi* epimastigotes undergoing mitosis and cytokinesis using iU-ExM (Figure 5). This approach enabled the visualization of mitotic spindle microtubules (MSTs), as well as the organization of the subpellicular microtubule network and the duplication of specialized cytoskeletal structures during cell division.

**Figure 5:**
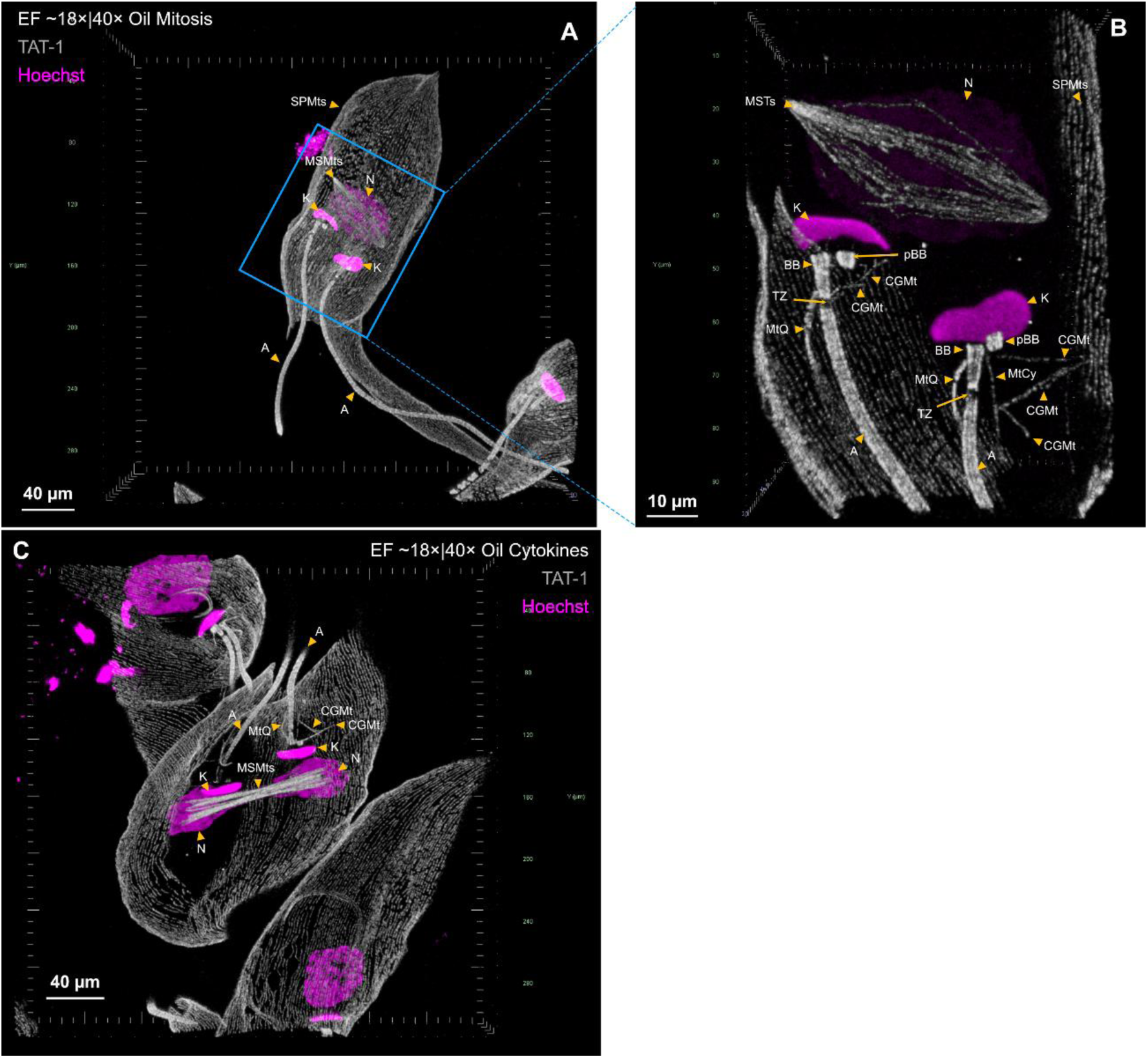
Visualization of mitotic spindle microtubules and cytoskeletal organization during mitosis and cytokinesis in *Trypanosoma cruzi* using iU-ExM. Representative confocal images of epimastigotes subjected to iterative ultrastructure expansion microscopy, achieving a final linear expansion factor of approximately 18x. Microtubules were labeled with the anti-TAT-1 antibody and a fluorescent secondary antibody (gray), while DNA was stained with Hoechst 33342 (magenta). (A) Confocal z-stack projection of an epimastigote undergoing mitosis, showing the subpellicular microtubule network (SPMTs), mitotic spindle microtubules (MSTs), nucleus (N), kinetoplasts (K), and axonemes (A). The blue rectangle indicates the region selected for detailed examination. (B) Projection generated from selected optical sections of the z-stack, highlighting the mitotic spindle and duplicated cytoskeletal structures, including the basal body (BB), probasal body (pBB), transition zones (TZ), microtubule quartets (MtQ), cytostome–cytopharynx-associated microtubules (MtCy), and Golgi complex-associated microtubules (CGMt). (C) Confocal z-stack projection of an epimastigote undergoing cytokinesis, showing the reorganization of the subpellicular microtubule network, mitotic spindle microtubules, and spatial segregation of nuclear chromatin and kinetoplasts during cell division. Scale bars: 40 µm (A, C) and 10 µm (B), corresponding to expanded sample dimensions. All images were acquired using a 40x oil-immersion objective.

In mitotic cells, confocal images revealed two kinetoplasts and a large nucleus, together with a prominent arrangement of microtubules corresponding to the mitotic spindle (Figure 5A). Projections generated from selected optical sections of the z-stack allowed a more detailed examination of the mitotic spindle and duplicated structures associated with the flagellar apparatus (Figure 5B). These included the basal and probasal body regions, transition zones, microtubule quartets, and microtubules associated with the Golgi complex. Notably, cytopharynx-associated microtubules were no longer detectable in mitotic parasites, consistent with the disassembly of the cytopharyngeal microtubule array during this stage of the cell cycle.

In cells undergoing cytokinesis, iU-ExM revealed the reorganization of the subpellicular microtubule network associated with daughter cell formation (Figure 5C). The images also showed the spatial separation of chromatin-containing nuclear regions and kinetoplasts, together with clearly distinguishable mitotic spindle microtubules. These observations allowed the organization of the cytoskeleton and the distribution of subcellular structures to be examined at different stages of cell division. The three-dimensional organization of these structures can be further appreciated in Supplementary Movie 3.

## 4. Discussion

In this study, we adapted iterative ultrastructure expansion microscopy (iU-ExM) for *Trypanosoma cruzi*, achieving a final linear expansion factor of approximately 18x. This approach enabled the acquisition of high-resolution confocal images of the parasite’s microtubule cytoskeleton, revealing the organization of both subpellicular microtubule arrays and specialized internal microtubule structures. Notably, iU-ExM also allowed for the detailed examination of cytoskeletal structures during mitosis and cytokinesis, providing new insights into the architectural remodeling associated with the parasite’s cell division.

Expansion microscopy has substantially enhanced the resolution capacity of conventional fluorescence microscopy by physically separating cellular structures within an expandable polymer network (Chen et al., 2015). Subsequent advancements, including ultrastructure expansion microscopy (U-ExM), have further improved the preservation and visualization of subcellular architecture (Gambarotto et al., 2019). In kinetoplastid parasites, Gorilak et al., 2021 demonstrated the utility of expansion microscopy for quantitative investigations of cytoskeletal organization, while Alonso, 2022 established U-ExM in *T. cruzi*, enabling the localization of various tubulin isoforms and isotypes. Building upon these developments and the iterative expansion strategy described by Louvel et al., 2023, our study extends the application of expansion microscopy to *T. cruzi*, achieving a significantly greater degree of physical expansion and improving the visualization of densely packed cytoskeletal structures.

One of the primary technical challenges in adapting iU-ExM for *T. cruzi* was the preservation and homogenization of its highly organized microtubule cytoskeleton. Unlike many mammalian cells, *T. cruzi* possesses a dense network of subpellicular microtubules that are closely associated with the plasma membrane, which contributes to maintaining cell shape and structural integrity (Vidal & De Souza, 2022). This organization poses a specific challenge for physical expansion, as insufficient homogenization can restrict the separation of adjacent structures. In our protocol, denaturation at 85 °C for 1h 30 minutes, combined with the use of a DHEBA-crosslinked hydrogel, proved compatible with subsequent expansion and the preservation of recognizable cytoskeletal architecture. This condition was essential for performing successive rounds of expansion while maintaining sufficient structural integrity for confocal microscopy imaging.

Another relevant aspect of the adapted protocol was the incorporation of immunostaining between the two expansion cycles. After the first expansion, the hydrogel was contracted in PBS to facilitate sample handling and incubation with antibodies. Immunostaining at this intermediate stage allowed for the labeling of microtubules prior to the second hydrogel embedding and subsequent expansion. This strategy follows the principle of iterative ultrastructure expansion microscopy, as described by Louvel et al., 2023, in which successive steps of hydrogel embedding enable further physical separation of the labeled structures. As reported by Louvel et al. 2023, post-expansion immunostaining often yields weak, inconsistent, or undetectable signals owing to restricted antibody diffusion within the dense iterative matrix. In our adaptation, the intermediate labeling procedure was compatible with the preservation of fluorescence (using an anti-TAT-1 antibody) during the second expansion, allowing for the visualization of microtubule arrangements in the final expanded specimens. Furthermore, performing immunostaining prior to the final expansion avoids the practical difficulties associated with antibody incubation in significantly larger hydrogel specimens.

The increased expansion factor was particularly advantageous for examining the subpellicular microtubule network, one of the most distinctive features of kinetoplastid cell architecture. Although this network has been extensively characterized by electron microscopy, its dense organization makes individual microtubules difficult to distinguish using conventional fluorescence microscopy. Following iU-ExM, the increased physical separation between neighboring microtubules enabled a more detailed visualization of their arrangement along the parasite cell body. Three-dimensional reconstructions further illustrated the spatial organization of the subpellicular network and highlighted the region associated with the cytostome, where the continuous subpellicular microtubule array is interrupted. These observations demonstrate the potential of iU-ExM for examining microtubule organization in three dimensions using confocal microscopy.

Beyond the subpellicular network, iU-ExM improved the visualization of specialized internal microtubule arrays associated with essential cellular structures. These included the microtubule quartet, cytostome–cytopharynx-associated microtubules, Golgi-associated microtubules, and structures associated with the basal and probasal bodies and the flagellar apparatus. The ability to distinguish these microtubule populations within the same expanded specimen provides valuable spatial information regarding their organization and relationships within the parasite. Previous studies using electron microscopy and expansion microscopy have demonstrated the complexity of specialized cytoskeletal structures in kinetoplastids, including those associated with the flagellar pocket and its biogenesis (Alcantara et al., 2014; Zelená et al., 2025). Our findings complement these approaches by enabling visualization of multiple microtubule arrays in the context of the overall cellular architecture of *T. cruzi*.

The application of iU-ExM further revealed distinct patterns of cytoskeletal organization associated with mitosis and cytokinesis in *T. cruzi* epimastigotes. In mitotic cells, the enhanced structural detail enabled the identification of mitotic spindle microtubules and duplicated components of the flagellar apparatus, highlighting the extensive cytoskeletal reorganization accompanying cell division. These findings are consistent with and further support previous ultrastructural observations reported by de Souza & Meyer, 1974 and Solari, 1980 using transmission electron microscopy, providing complementary evidence of the cytoskeletal rearrangements associated with mitosis in *T. cruzi*.

Notably, cytopharynx-associated microtubules were not detected at this stage, consistent with previous ultrastructural observations demonstrating the disassembly of the cytostome–cytopharynx complex during the cell cycle (Alcantara et al., 2017). In cells undergoing cytokinesis, iU-ExM provided a detailed view of the subpellicular microtubule network and its organization in relation to the segregating kinetoplasts and nuclear chromatin, together with the microtubules of the mitotic spindle. These observations highlight the spatial coordination of specialized microtubule arrays and nuclear and kinetoplast segregation during parasite division. Although derived from fixed specimens, these images provide structural snapshots of successive stages of the cell cycle, demonstrating the potential of iU-ExM to investigate cytoskeletal remodeling and the spatial relationships between microtubule structures and organelles during *T. cruzi* cell division.

Despite these advantages, some limitations should be considered. The effective resolution achieved by expansion microscopy depends not only on the expansion factor but also on the optical resolution of the microscope, the labeling strategy, and the structural preservation of the specimen. Therefore, the approximately 18x expansion achieved in this study should not be interpreted as direct evidence of a particular absolute spatial resolution without additional quantitative validation. Similarly, although the preservation of recognizable cytoskeletal structures supports the applicability of the protocol, systematic measurements would be required to establish the degree of local expansion isotropy and potential structural distortions. The combination of iU-ExM with quantitative image analysis and complementary electron microscopy could help address these questions and further establish its suitability for precise structural measurements.

Overall, our findings establish iU-ExM as a valuable approach for investigating the cytoskeletal and subcellular architecture of *T. cruzi*. By combining successive physical expansion with fluorescence labeling and confocal microscopy, this methodology enables detailed visualization of microtubule arrangements that are difficult to resolve using conventional fluorescence imaging. Its application to parasites undergoing mitosis and cytokinesis further highlights its potential for investigating the structural organization and remodeling of the kinetoplastid cytoskeleton throughout the cell cycle.

## Acknowledgments

We would like to thank Verônica dos Santos and Paloma Lira de Oliveira Chagas for their valuable technical assistance and support throughout this study.

## Author Contributions

**Normanda Souza-Melo:** conceptualization, data curation, formal analysis, investigation, visualization, methodology, validation, writing – original draft. **Giovanna Henriques de Souza:** investigation, visualization. **Sharmila Ortiz:** methodology, data curation, and visualization. **Wanderley de Souza:** conceptualization, resources, data curation, supervision, writing – review and editing.

---

**Supplementary Figure S1:**
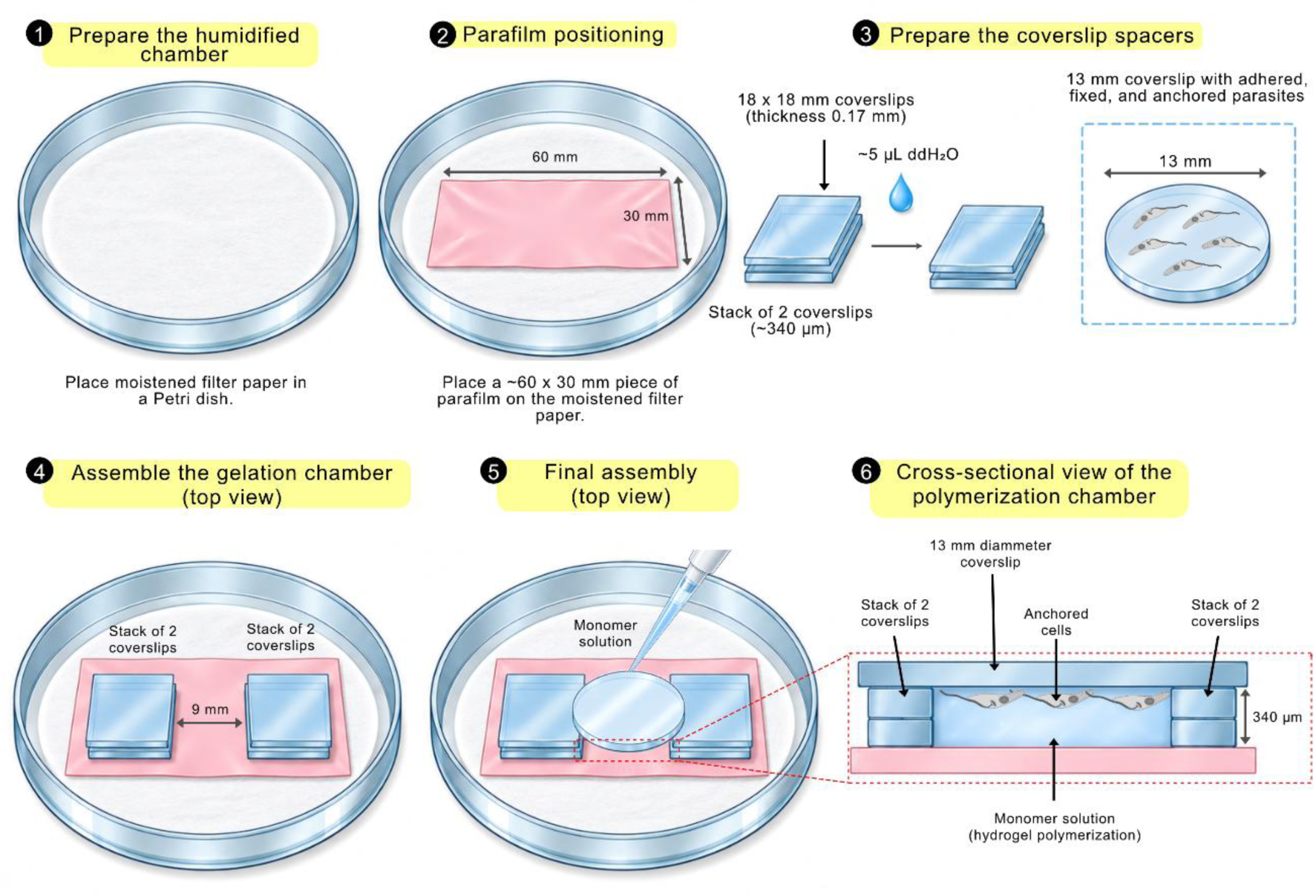
Schematic representation of the gelation chamber assembly for iterative ultrastructure expansion microscopy (iU-ExM) in *Trypanosoma cruzi*. The diagram illustrates the preparation and assembly of the gelation chamber used for the first hydrogel polymerization step. (1) A humidified chamber was prepared using a Petri dish containing water-moistened filter paper. (2) A piece of Parafilm (approximately 60 x 30 mm) was placed over the moistened filter paper. (3) Two spacers were prepared, each consisting of two stacked 18 x 18 mm glass coverslips (0.17 mm thick each), held together with approximately 5 µL of ultrapure water. A 13-mm round coverslip containing previously adhered, fixed, and anchored *T. cruzi* epimastigotes was prepared for subsequent placement over the spacers. (4) The two coverslip stacks were positioned parallel on the Parafilm, approximately 9 mm apart, allowing the 13-mm round coverslip to rest on both spacers. (5) The 13-mm coverslip was placed over the two spacers, with the parasites facing the Parafilm, and the hydrogel monomer solution was introduced into the space between the coverslip and the Parafilm. (6) Cross-sectional view of the final assembly, showing how the two coverslip stacks serve as spacers, creating a polymerization chamber approximately 340 µm thick between the round coverslip and the Parafilm.

**Supplementary Movie 1: Three-dimensional visualization of the subpellicular microtubule network in *Trypanosoma cruzi* epimastigotes following iU-ExM.** Three-dimensional visualization generated from confocal z-stack images of a *T. cruzi* epimastigote subjected to iterative ultrastructure expansion microscopy (iU-ExM), achieving a final linear expansion factor of approximately 18x. Microtubules were immunolabeled with the anti-TAT-1 antibody (gray). The reconstruction highlights the highly organized subpellicular microtubule network along the parasite cell body and the region corresponding to the cytostome, where this network is interrupted. The movie complements Figure 4A by illustrating the spatial organization of the subpellicular microtubules.

**Supplementary Movie 2: Three-dimensional visualization of specialized internal microtubule arrays in *Trypanosoma cruzi* following iU-ExM.** Three-dimensional visualization generated from confocal z-stack images of a *T. cruzi* epimastigote following approximately 18x expansion by iU-ExM. Microtubules were immunolabeled with the anti-TAT-1 antibody (gray). The movie highlights the spatial organization of specialized internal microtubule arrays, including the microtubule quartet (MtQ), cytostome–cytopharynx-associated microtubules (MtCy), Golgi complex-associated microtubules (CGMt), and structures associated with the flagellar apparatus. This visualization complements Figure 4C by providing additional information on the three-dimensional arrangement and spatial relationships of these structures.

**Supplementary Movie 3: Three-dimensional visualization of cytoskeletal organization during cell division in *Trypanosoma cruzi* following iU-ExM.** Three-dimensional visualization generated from confocal z-stack images of a *T. cruzi* epimastigote undergoing cytokinesis following approximately 18x expansion by iU-ExM. Microtubules were immunolabeled with the anti-TAT-1 antibody (gray), and DNA was stained with Hoechst 33342 (magenta). The movie illustrates the organization of the subpellicular microtubule network, mitotic spindle microtubules (MSTs), and the spatial distribution of nuclear chromatin and kinetoplasts during cell division. This reconstruction complements Figure 5C by providing a three-dimensional view of cytoskeletal architecture during cytokinesis.

